# Exploring the Genomic Landscape of the GP63 family in *Trypanosoma cruzi*: Evolutionary Dynamics and Functional Peculiarities

**DOI:** 10.1101/2024.02.17.580826

**Authors:** Luisa Berná, María Laura Chiribao, Sebastian Pita, Fernando Alvarez-Valin, Adriana Parodi-Talice

**Affiliations:** Laboratorio de Interacciones Hospedero–Patógeno—UBM, Institut Pasteur Montevideo, Montevideo 11400, Uruguay; Laboratorio de Genómica Evolutiva, Facultad de Ciencias, Universidad de la República, Montevideo 11400, Uruguay; Departamento de Bioquímica, Facultad de Medicina, Universidad de la República, Montevideo 11800, Uruguay; Sección Genética Evolutiva, Instituto de Biología, Facultad de Ciencias, Universidad de la República, Montevideo 11400, Uruguay

**Keywords:** Zinc-metalloprotease GP63, Leishmanolysin, *Trypanosoma cruzi*, multigene family evolution

## Abstract

We analyzed the complete set of GP63 sequences from the parasitic protozoa *Trypanosoma cruzi*. Our analysis allowed us to refine annotation of sequences previously identified as functional and pseudogenes. Concerning the latter, we unified pseudogenic fragments derived from the same functional gene and excluded sequences incorrectly annotated as GP63 pseudogenes. We were able to identify eleven GP63 gene groups, which are sharply defined and have a high intra-group sequence identity. The sequences of each group showed a strong preference for genomic compartments. Some groups are located in the core and others in disruptive compartments of the *T. cruzi* genome. Groups located in the core compartment often contain tandem arrays of GP63 genes. On the contrary, genes from groups located in the disruptive compartment tend to be surrounded by genes encoding surface proteins such as MASP, mucins and trans-sialidases. Analysis of the immediate GP63 environments showed differences that may be the result of different genomic dynamics in these two compartments. Interestingly, each GP63 group showed a particular mRNA expression profile and some groups contain members that are differentially expressed between life cycle stages, being expressed at higher levels in trypomastigotes than in the replicative forms. This suggests that these groups of GP63 proteins may play a relevant role in the infective stage. The analysis of the M8 domain, that defines the GP63 protein family, allowed us to recognize that each group presented peculiarities in the conserved sites as well as in the presence of the predicted signal peptide and GPI anchor site. Phylogenetic analysis of the GP63 sequences, including other species of the genus *Trypanosoma* as well as other kinetoplastids, showed that ten of the 11 groups of *T. cruzi* not only are also present in the other *Trypanosoma* species but also are exclusive of genus, suggesting that the diversification of these subfamilies took place before speciation. However, each species then followed a different evolutionary path, amplifying specific groups in unique ways.

**Data summary:** The authors confirm all supporting data, code and protocols have been provided within the article or through supplementary data files.

**Impact statement:** Our study contributes to the understanding of the GP63 gene family in *Trypanosoma cruzi*, a crucial protein for the parasite’s infectivity and evolution. We refined the annotation of GP63 sequences, identifying eleven distinct gene groups with distinctive preferences for genomic compartments -some in the core, others in the disruptive compartment. This distribution hints at varied genomic dynamics and potential roles in the parasite’s life cycle, especially since some groups show enhanced expression in infective stages, suggesting their importance in disease transmission.

Our exploration into the GP63 sequences’ M8 domain revealed group-specific peculiarities in conserved sites and structural motifs, emphasizing functional diversity. Phylogenetic analysis across *Trypanosoma* species highlighted the evolutionary uniqueness of these gene subfamilies within the genus, underscoring their role in the species’ distinct evolutionary paths and amplification patterns.

## INTRODUCTION

Chagas disease, caused by the *Trypanosoma cruzi* parasite, affects approximately 7 million individuals worldwide and is classified as a neglected disease by the World Health Organization (WHO Chagas disease). Primarily prevalent in rural regions of Latin America, this disease holds a significant impact. The progression of Chagas disease heavily relies on the interaction of parasitic virulence factors and the defensive responses of the mammalian host. In this context, identifying parasite molecules that engage with the host’s immune or signaling systems could present novel targets crucial for disease intervention.

The surface of the parasite, especially during its infective stages, serves as a crucial interface for interaction with the vertebrate host. Surface proteins belonging to families such as mucins, MASPs and trans-sialidases are prominently expressed in this domain. While advancements have been achieved in understanding the surface architecture of these parasites (1–3), the precise roles of individual components in host-parasite interactions, as well as the dynamic expression patterns of all constituents during the infection process, remain elusive.

Although different species exhibit distinct types of surface components, GP63 glycoproteins are the only surface protein components that are universally present across all trypanosomatid species. The genes encoding GP63 proteins constitute a family that exhibits considerable variability in both number and structure across these species (4).

The GP63 protein is a metalloprotease that belongs to the group of zinc-dependent endopeptidases. Sequences akin to GP63 are also found in other insect and plant parasites (4). The diversification of trypanosomatid genomes has been accompanied by the rapid evolution of their multigene families, including the GP63 family, which underwent substantial modification after the emergence of parasitism. This is particularly evident when comparing these genomes to those of free-living kinetoplastids (5). Phylogenetic studies of the GP63 family showed that both *T. cruzi* and *Leishmania* sp. have independently expanded their gene repertoires (5). Therefore, this family has had a long evolutionary history, and it has been postulated that co-evolutionary processes of parasites and their hosts have significantly influenced the structure and functionality of its members (4,6).

The GP63 protein was first described in *Leishmania* (7), where it has been extensively studied ever since. Some members are known to be expressed on the surface, others are secreted, and others remain intracellular (8). It is the most abundant protein in the parasite (500,000 molecules/parasite). GP63 in *Leishmania* plays several roles relevant for infection such as: inhibition of complement-mediated lysis, facilitation of phagocytosis, degradation of extracellular matrix components, alteration of host cell signaling pathways and immune evasion (8–10). This multiplicity of functions suggests a preponderant role for parasite biology. In contrast to the extensive evidence obtained in *Leishmania*, very little is known about the functions of this family in *T. cruzi*, despite its abundance in the genome.

Some studies showed that members of the GP63 family of *T. cruzi* may be involved in infection. This is supported by the fact that pre-incubation with specific antibodies partially blocked infection of Vero cells by trypomastigotes (11). Other studies showed that GP63 RNAs are differentially expressed depending on the stage of the parasite life cycle (12). A few studies on metalloproteinase activity detected on gelatin SDS-PAGE gels have been reported for a group of GP63 proteins of 75-78 kDa (11,13)

The variability in the GP63 gene family across *Leishmania* species, ranging from 4 to 33 genes, contrasts starkly with the findings in *Trypanosoma cruzi*. The hybrid strain CLBrener’s genome, initially reported in 2005 by El-Sayed et al.(14), contained 425 GP63 copies. However, more recent genomic analyses using long-read, third-generation sequencing techniques have provided a more accurate assembly and annotation, revealing 378 GP63-encoding genes in the Dm28c strain of *T. cruzi* and 718 in the hybrid strain TCC (15). This sequencing approach not only offered a more realistic gene count by resolving previously collapsed repetitive regions but also unveiled a compartmentalized, or ‘bipartite’, genome structure with distinct GC content. Specifically, we described the *T. cruzi* genome as comprising of a core compartment, which contains genes conserved across trypanosomatids and encodes both known and hypothetical proteins, and a disruptive compartment. This latter is composed by genomic regions that are unique to *T. cruzi* and enriched with species-specific gene families such as mucins, mucin-associated surface proteins (MASPs), and trans-sialidases. Notably, GP63 family members are distributed across both genomic compartments. The specific attributes of the genes within these compartments, reflecting their unique evolutionary and functional dynamics, are yet to be fully understood.

In *Leishmania* species, at least 60% of the amino acids in the GP63 protein, also known as Leishmanolysin, are conserved. This includes the short zinc-binding motif HEXXH and 19 cysteine residues distributed along the sequence. Following synthesis, Leishmanolysin undergoes several post-translational modifications: the signal sequence is cleaved in the endoplasmic reticulum, N-glycosylation is applied, and roughly 25 C-terminal amino acids are substituted by a GPI anchor. The protein is further processed by proteolytic cleavage of a propeptide, yielding the mature form. The pro-peptide sequence harbors a cysteine residue conserved across all GP63 variants in different species (16). In *Leishmania major*, this cysteine residue is implicated in a ‘cysteine switch’ mechanism, where it plays a role in inhibiting protease activity by chelating the zinc atom at the active site. The mature protein in *L. major* begins at Val100, features three N-glycosylation sites at Asn300, Asn407, and Asn534, and concludes at Asn577, which is covalently attached via ethanolamine to the carbohydrate portion of the GPI anchor.

The zinc-binding motif HEXXH located in the amino-terminal domain of GP63 is the characteristic site of metalloproteases. Research demonstrates that the two histidines and one glutamic acid within this motif are conserved across all GP63 sequences in the trypanosomatids studied, playing a crucial role in proteolytic activity (17). Mutations in any of these three residues result in a complete loss of catalytic function. Specifically, a mutation from glutamic acid to aspartic acid at position E165 in the GP63 of *L. major* leads to a loss of protease activity (18).

In this research, we aimed to extensively characterize the GP63 gene family in the genome of *T. cruzi*, examining its subtypes, domains, genomic organization, evolutionary dynamics, expression profiles, and structural characteristics. Through this detailed analysis, we seek to contribute with enough knowledge necessary to elucidate the unique role of these proteins to pathogenesis and immune system evasion.

## RESULTS AND DISCUSSION

### 1 The metalloprotease GP63 family contains multiple functional gene copies and several pseudogenes in the genome of *Trypanosoma cruzi*

To analyze the GP63 family of metalloproteases, we utilized the annotated Dm28c genome, sequenced using long-read technology (15). This approach is crucial for including all family copies. In this genome, we identified 96 complete gene sequences encoding GP63 proteins. Furthermore, the genome possesses annotations for 281 potential GP63 pseudogenes. The primary criterion for classifying a sequence as a pseudogene was its length; specifically, sequences encoding polypeptides of 460 or fewer amino acids were deemed pseudogenes.

Upon examining the length distribution of the entire sequence population, a broad range of variation is evident, displaying a clear division between potential functional sequences and pseudogenes (Figure 1A). Furthermore, sequences classified as functional also exhibit significant variability. In fact, the distribution is bimodal, peaking at lengths of approximately 560 and 700 amino acids (Figure 1B).

**Figure 1:**
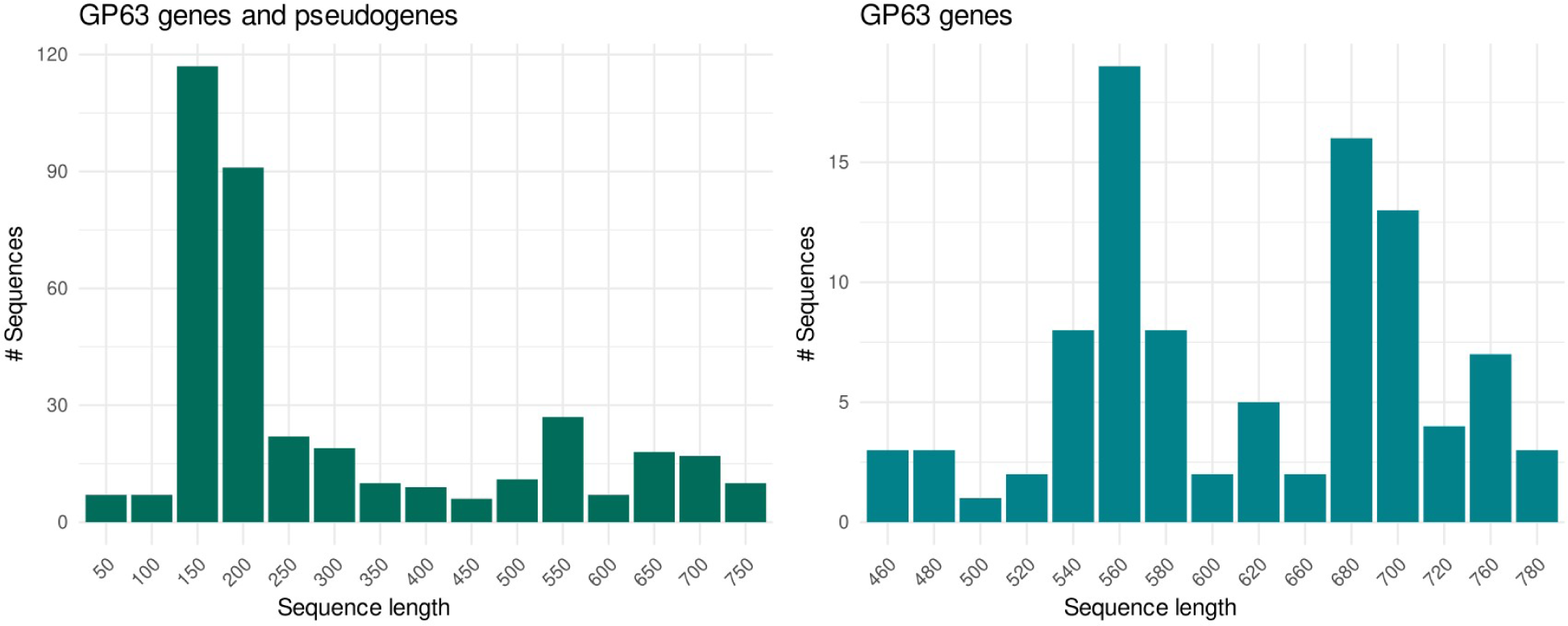
Distribution of GP63amino acid sequence lengths in *T. cruzi* Dm28c. This figure displays two distinct length distributions: on the right, the combined length distribution of both GP63 genes and pseudogenes; on the left, solely GP63 functional genes.

Using length as the only criterion to categorize sequences as pseudogenes or functional, is incomplete to get a full grasp. These aspects led us to refine the annotation by incorporating additional criteria. Specifically, to classify sequences as functional, we now consider a sequence as such if it is longer than 460 amino acids, contains a full-length peptidase M8 domain and at least in the majority of the cases, presents a signal peptide (refer to Section 4 for further details).

Concerning pseudogenes, it is important to bear in mind that many sequences classified as pseudogenes may actually represent remnant fragments of the same former functional GP63 gene that underwent pseudogenization, as already reported for most trypanosomatid genes (19). In fact, among the 281 sequences previously classified as pseudogenes, 63 are observed in pairs or groups, either juxtaposed, overlapping, or separated by short distances. Analysis of these sequences shows that they are indeed sister fragments that originated by nonsense mutations or indels producing frameshifts of a previously intact coding sequence. Segments belonging to these groups are now classified as being parts of the same pseudogene. We also found that 29 pseudogenic fragments are not really related to GP63. In all likelihood, these fragments were originally incorrectly named as GP63 because of flawed transference of information based on homology. The revised list, now comprising 185 pseudogenes, is presented in Supplementary Table 1.

It is worth noting that many of these pseudogenic fragments retain high DNA sequence identity with complete (functional) genes, suggesting their status as “young” pseudogenes, namely nonsense mutations or indels causing pseudogenization occurred relatively recently. Moreover, comparative analysis of paired pseudogenic fragments, particularly at the indels sites, shows that several pairs (or groups) contain mutations at identical positions. This pattern suggests that they are derived from duplication events that have occurred after pseudogenization (Supplementary Figure 1).

### 2 Eleven groups of GP63 genes are present in *Trypanosoma cruzi* genome

Previously, Cuevas et al. (11) conducted PCR analyses and identified *T. cruzi* sequences homologous to GP63 genes from *Leishmania*. They categorized GP63 genes in *T. cruzi* into three groups: Tcgp63-I and Tcgp63-II, and Tcgp63-III, consisting of pseudogenes. Subsequent re-analysis of the *T. cruzi* GP63 family genes led to their classification into four distinct groups (4). Within each group, the sequences exhibit 80% similarity, whereas between groups, the similarity drops to 30%. These authors identified a conserved segment of 100 amino acids following the active site, shared across all sequences, likely corresponding to the M8 domain, a feature ubiquitous to all GP63 sequences.

In this study, thanks to the development of third-generation technologies that allow sequencing of increasingly better-quality long reads, new *T. cruzi* genomes, with higher accuracy have become available. This advancement enabled us to conduct an exhaustive comparative analysis of the 96 annotated GP63 gene sequences, using both nucleotide and protein sequences (Supplementary Table 2). This approach led to the identification of 11 distinct groups within the GP63 gene family. Each group has a characteristic average length of sequences, which in most groups has little variation (Supplementary Figure 2). This classification is evident in a phylogenetic analysis performed with the complete amino acid sequence (Figure 2A, Supplementary Figure 3). Furthermore, an amino acid identity matrix, created through all-against-all pairwise comparisons, corroborated the observed clustering patterns (Figure 1B, Supplementary Table 3). This matrix was visualized using Gephi, where each node represents a GP63 sequence and the color of each edge denotes the level of identity, and an edge weight threshold of 65% was applied (Figure 2B).

**Figure 2:**
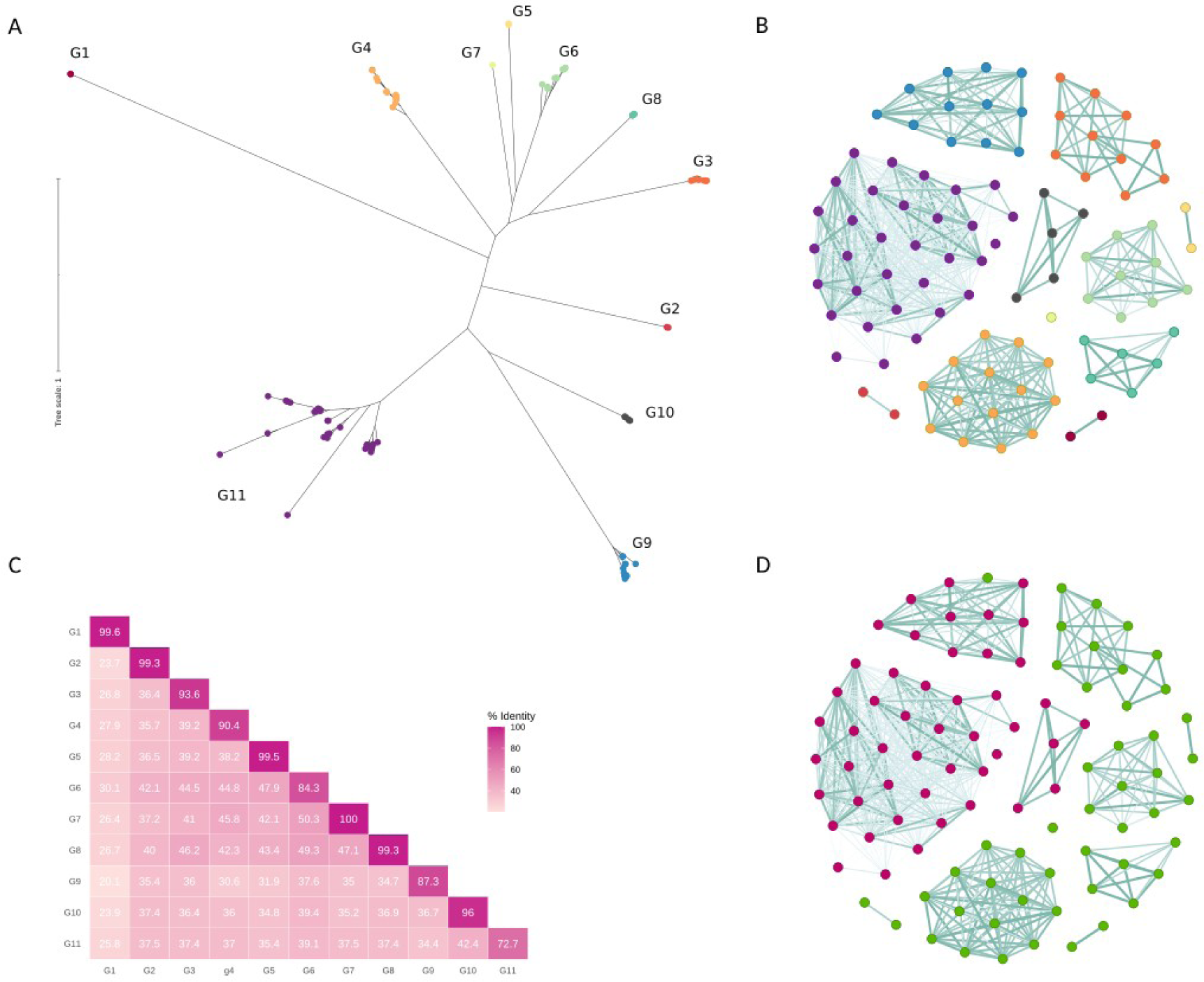
Distinct Clustering of GP63into 11 Groups. A) Phylogenetic analysis of amino acid sequences of the 96 GP63 genes from *T. cruzi* Dm28c, illustrating the division into 11 unique groups. B) Network visualization in Gephi of the pairwise identity distances among the 96 amino acid sequences of GP63 genes, where nodes symbolize genes (color coding matches the phylogeny in A), and the color of each edge indicates the level of identity. C) Identity matrix displaying average intra-group and inter-group amino acid identities, highlighting the distinctiveness of each group. D) Network visualization similar to B, with genes colored according to the genomic compartment to which they belong: green for the core compartment and pink for the disruptive compartment.

Additionally, the identity matrix was condensed into a matrix representing average intra-group and inter-group identity, as depicted in Figure 2C. The intra-group amino acid identity levels are predominantly homogeneous, exceeding 90% in 8 out of 11 groups. In particular, three groups showed more variability, especially group G11, which is the largest and exhibited the lowest homogeneity with 73% mean identity. In contrast, amino acid identities between groups were notably lower, never exceeding 48%, indicating significant divergence among groups. This pronounced divergence between groups, together with the strong homogeneity within most groups, justifies their clustering into 11 distinct groups, as highlighted in Figure 2. Further expanding on this classification, our BLAST analysis of *T. cruzi* genes and pseudogenes enabled us to assign each pseudogene a parental group, thereby extending our clustering analysis beyond genes to include pseudogenes. As revealed in Supplementary Table 2, a significant proportion of these pseudogene sequences predominantly correspond to groups G9 and G11, both of which are located in the disruptive compartment. This inclusion of pseudogenes in our analysis underscores the comprehensive nature of the classification, affirming the robustness of the 11-group clustering in capturing the genetic diversity within *T. cruzi*.

### 3 Non-random distribution of GP63 gene groups within the *T. cruzi* genome

The genome of *T. cruzi* is composed of two clearly defined compartments, the so-called disruptive and core compartments (15). These genomic compartments are recognized for their distinct base composition (% GC), gene content, and likely exhibit differential regulation and functions within the parasite, alongside unique evolutionary trajectories (20). The core compartment contains genes conserved between different *T. cruzi* strains but also largely shares synteny conservation with other trypanosomatids. On the other hand, the disruptive compartment is composed almost exclusively of multigene families for surface proteins. While the majority of gene families encoding surface proteins are situated within the disruptive compartment, GP63 does not follow this pattern. Our findings reveal that the GP63 gene family is evenly distributed across both compartments (Figure 2D and Supplementary Figure 4). This contrasts with *Leishmania major*, which harbors a single tandem array with six GP63 genes in its genome (21). Unlike *L. major*, *T. cruzi* shows a wide dispersion of GP63 genes throughout its genome. Furthermore, the genomic positioning of the various GP63 groups shows a strong correlation with these specific genomic regions. Notably, genes belonging to groups G1 to G8 are all found in the core compartment, whereas genes from groups G9, G10 and G11 are present almost exclusively in the disruptive compartment (Figure 2D and Supplementary Figure 4).

We also performed a local gene environment analysis aiming to identify the most frequent neighbors of GP63 genes. The three most abundant genes were identified at both, 3’ or 5’ neighborhood of each GP63 coding sequence. By neighborhood we understand the three closest neighbors. We found that depending on whether the GP63 gene is located in the core or disruptive compartment, its gene environment is quite different. While genes in the core compartment mostly have a sire-like repeat sequence immediately upstream or downstream (Figure 3A), GP63 genes located in the disruptive compartment are found together with MASP, RHS and TS genes (Figure 3B and 3C). In addition, many GP63 genes are distributed in tandem repeats, and this is basically restricted to groups found in the core compartment (Figure 3C). In many cases, these tandem arrays do not contain exclusively the GP63 sequences but also the accompanying gene/s, meaning that the repetition unit can be somewhat larger.

**Figure 3:**
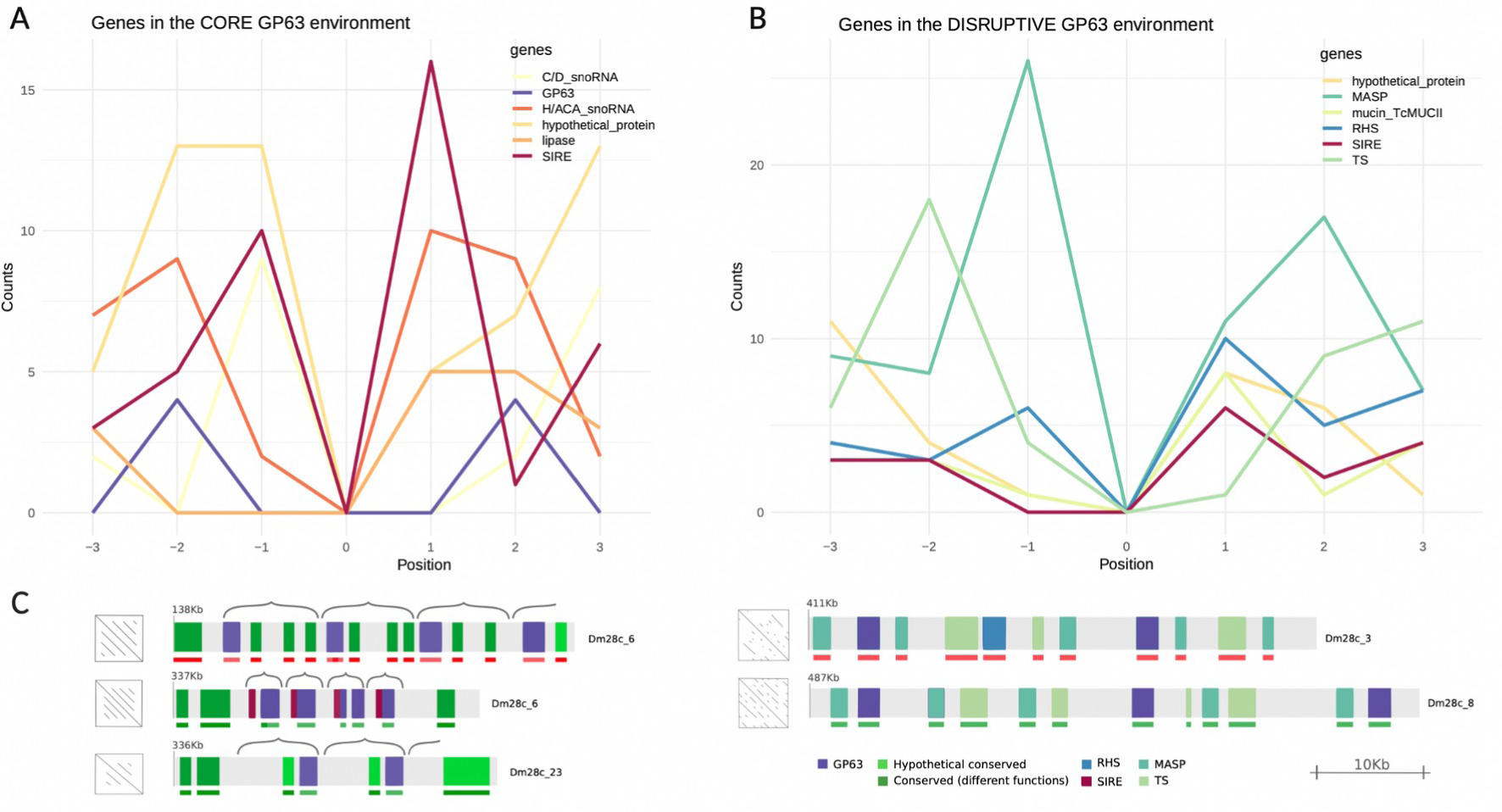
GP63 Gene Environment Composition. This figure illustrates the predominant genes found adjacent to each GP63 gene, focusing on the three genes upstream and downstream. A) Analyzes the environment of GP63 genes within the core compartment. B) Analyzes the environment of GP63 genes within the disruptive compartment. C) Provides examples of GP63 gene environments in both compartments, with visuals sourced from http://bioinformatica.fcien.edu.uy/cruzi/browser/dm28c.500.php. For these examples, the contig and starting location of the depicted fragments are specified. Each region presents a schematic dotplot, showing the organization of the tandem repeats.

The short interspersed repetitive element (SIRE)-like sequences, exclusive to *T. cruzi*, are prevalent throughout its genome, numbering between 1,500 and 3,000 copies (22). SIREs are often associated with protein-coding genes and are typically found in the 3’ untranslated regions (UTRs) of mRNAs, frequently contributing to polyadenylation sites (23). Moreover, SIRE elements are components of VIPER, a distinctive retroelement which is presumably active in *T. cruzi*. The presence of this type of sequences in the UTRs of several of the GP63 genes may have implications in the regulation of their expression. In fact, it is known that elements present in the UTRs influence their gene expression (24). Furthermore, the observed enrichment of SIRE elements in association with GP63 genes, especially within the core compartment, suggests their potential involvement in the gene duplication mechanisms of GP63 genes. Conversely, within the disruptive compartment, the genomic dynamics of GP63 genes are likely influenced by the amplification of other multigenic sequences, including mucins, RHS, trans-sialidases, and MASPs.

### 4 Signal and domain recognition in predicted GP63 amino acid sequences

According to the MEROPS database, the peptidases of the M8 family are zinc-metallopeptidases belonging to the MA clan of metallopeptidases, characterized by an active site containing a HEXXH motif involved in the coordination of the zinc ion required for catalytic activity. This clan includes several subfamilies and is known for its functional diversity, including peptidases involved in extracellular protein degradation, peptide signal processing, and other critical functions in both pathogens and their hosts. M8 family proteases are notably prevalent across various protozoan organisms, with *T. cruzi* boasting the largest collection of such proteins (298).

All identified sequences were found to contain a peptidase M8 domain (Leishmanolysin family, clan MA(M), InterPro Acc IPR001577). The Metzincin clan of metalloproteinases possess a consensus sequence that includes not only the two histidines that bind the zinc ion and the general-base glutamate (HEXXH), but also an extended motif with and additional strictly conserved glycine and a third zinc-binding histidine located further along the sequence. Metzincins are also characterized by a conserved methionine-containing 1,4-β-turn, known as the Met-turn (25). A distinctive aspect of leishmanolysins is the presence of a large insertion between the glycine and the third zinc-binding histidine. Our analysis focused on the presence of these conserved elements within the active site of the M8 domain. The zinc-binding site is preserved in most of the protein sequences, with extensive conservation observed beyond the second histidine, indicating a limited scope for variation in these positions, which consistently feature alanine and leucine (Figure 4). Intriguingly, two GP63 groups, G9 and G10, have a mutation in the highly conserved glutamic acid residue within the active site, by leucine and phenylalanine respectively. While the impact of this mutation on protein activity is unclear, it does not appear to influence gene expression. In fact, these genes exhibit RNAseq expression levels that are comparable to or exceed those of the other groups (see Figure 5). Furthermore, the variable number of residues between the conserved glycine in the active site and the third histidine is striking. While most of the groups have similar or exact spacing to the *Leishmania* sequences (62 amino acids), group G3 contains a significantly larger insertion of 150 to 270 residues. These insertions split the M8 domain into two segments, significantly increasing the distance between the third histidine and the preceding two histidines. The significance of this change in protein structure is not known but would merit further study in terms of its possible functional impact. In the case of group G1, this spacing is 71 residues. In addition, a Met-turn characterized by a highly conserved methionine is present in all groups except for this group G1. As illustrated in Figure 4, each group exhibits unique characteristics regarding these elements.

**Figure 4:**
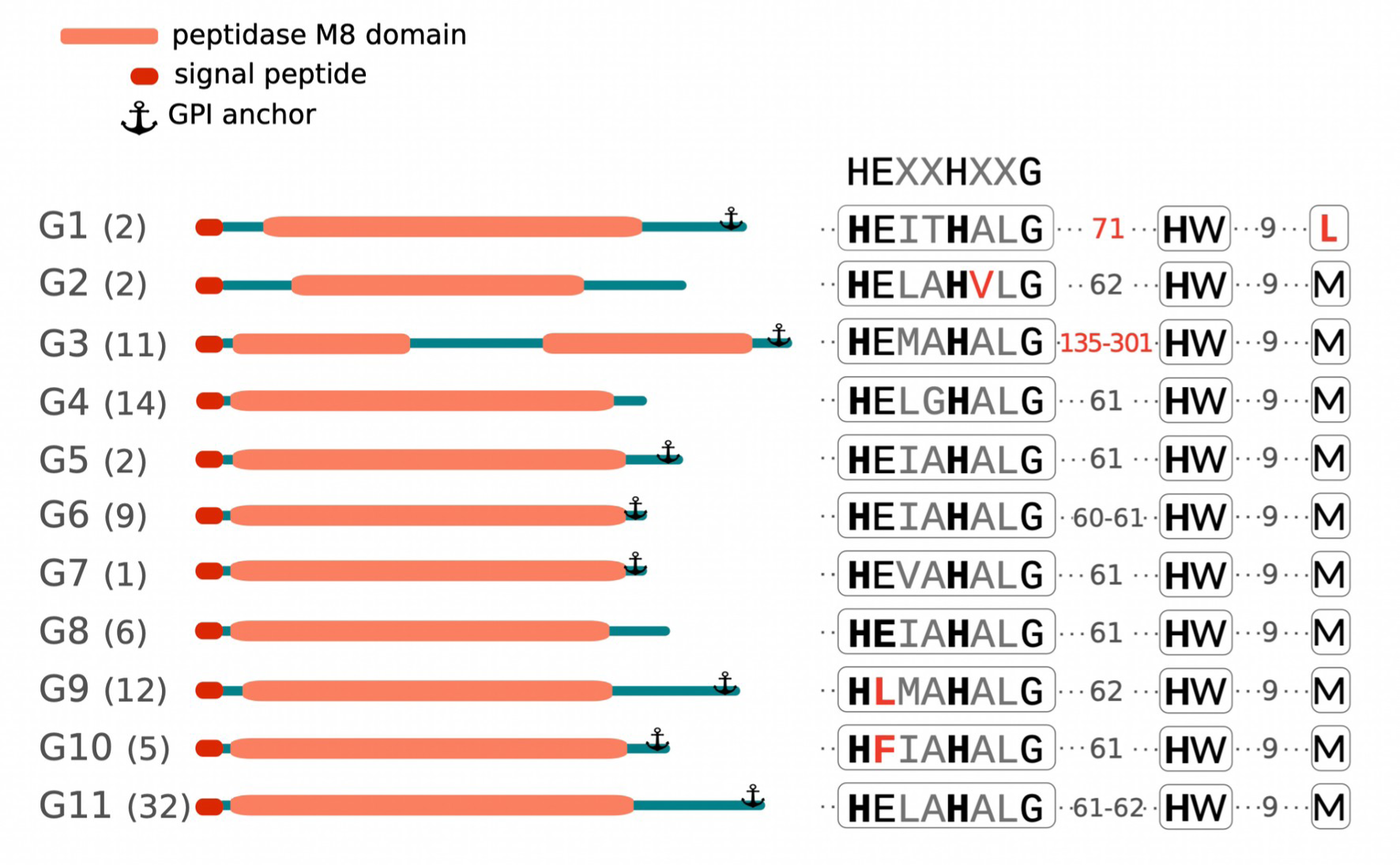
Structure of GP63 Group proteins. This figure provides a schematic overview of each GP63 group, depicting protein length, the location of the M8 domain, and the presence of signal peptides and GPI anchors. On the left side, each group is indicated and the number of sequences in each group is given in brackets. On the right side, the catalytic domain HEXXHXXG, the spacer to the subsequent histidine, and the conservation of the Met-turn are illustrated, along with all group-specific variations highlighted in red.

**Figure 5:**
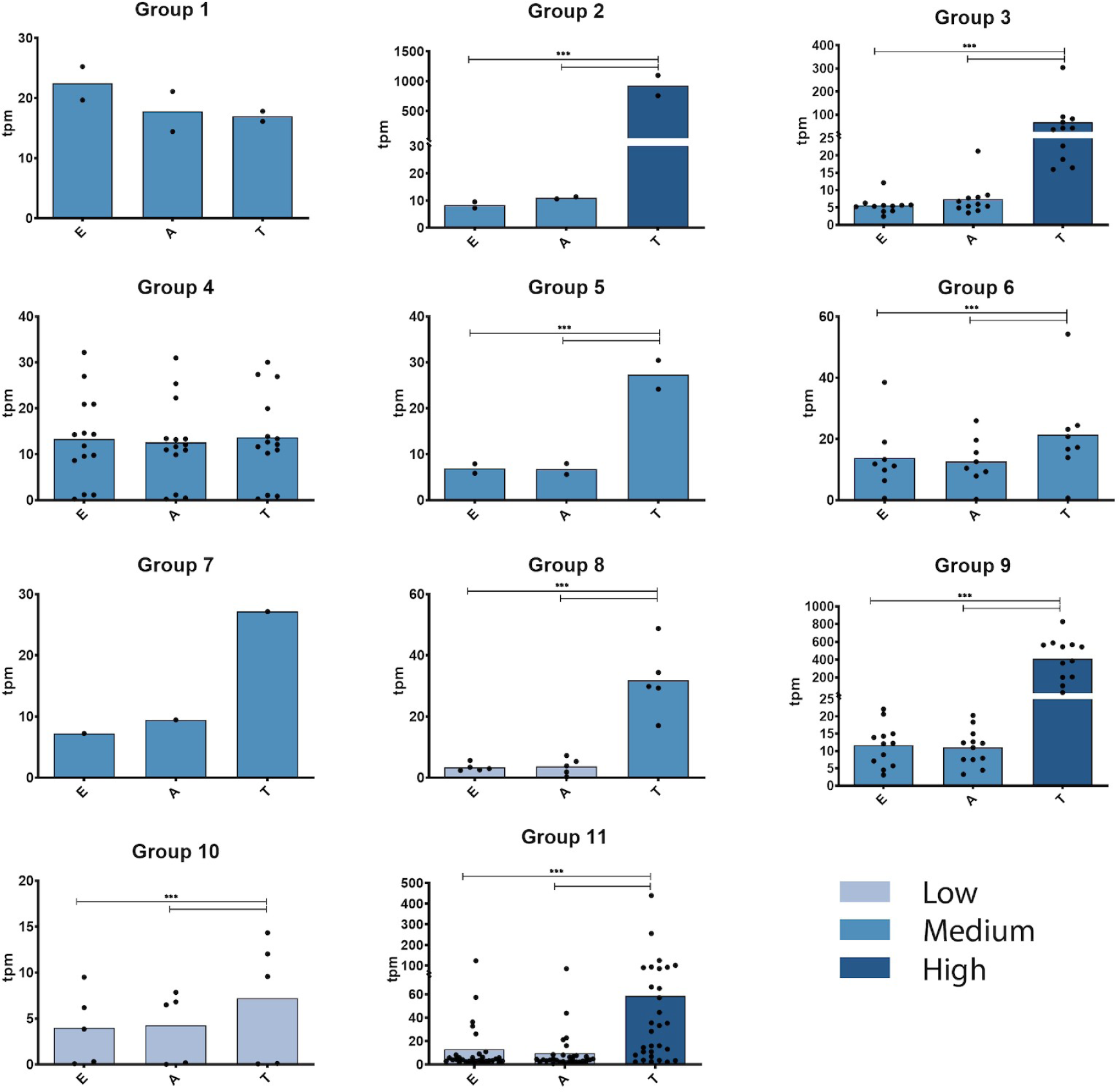
Expression profiles at the ARNm levels in the different GP63 groups. Each graph shows the expression profiles of the genes in each GP63 group. On the y-axis, expression levels are expressed in transcripts per million (tpm). Levels of expression were categorized as low, medium and high, and coloured differently. On the x-axis are the three stages of *T. cruzi*: Epimastigotes (E), Amastigotes (A) and Trypomastigotes (T). The black dots in each graph represent each individual gene. Statistical analysis of differential expression between stages was performed by 2 way ANOVA. *** represents a p-value <0.001

Mutations at H264, E265 and H268 of the active site of the *L. major* GP63 have been described to completely abrogate catalytic function (18). The presence of the essential active site elements in the majority of *T. cruzi* GP63 sequences suggests that these are proteins active in their protease function. In contrast, the absence of the highly conserved E residue of the active site of the sequences belonging to groups G9 and G10 of *T. cruzi*, which has been considered essential in the *L. major* protein, suggests that these two groups of sequences are not functional proteases. However, it should be noted that these proteins could have other functions, as has been described for the trans-sialidase family, a family that should be renamed because most of its members have no enzymatic activity due to a mutation in the active site (26). However, these proteins retain other important functions by facilitating interaction with host cell receptors, promoting signaling and cell invasion of trypomastigote forms (27).

The structural analysis of GP63 amino acid sequences revealed a predicted N-terminal signal sequence across almost all GP63 variants, ranging from 23 to 37 amino acids depending on the sequence. Only in seven sequences a signal peptide could not be predicted, due in almost all cases to N-terminal deletions. Of these, four sequences corresponded to group G11, one to G9, one to G8 and one to G4. Conversely, predictions for a GPI anchor site yielded inconsistent results. Most sequences from groups G1, G3, G5, G6, G7, G9, G10, and G11 feature a GPI anchor site, whereas sequences from groups G2, G4, and G8 appear to lack this site (Figure 4 and Supplementary Table 1). Furthermore, our examination of putative protein features, specifically the N-glycosylation sites across the 96 proteins, indicated a variability in the number of N-glycosylation sites among the groups. Notably, group G9 exhibited the highest count of glycosylation sites, as detailed in supplementary Figure 5.

The amino- and carboxyl-terminal ends of the GP63 sequences show the greatest variability between the different members of each group, both in terms of length and sequence composition. For leishmanolysin, as well as for several other families of metzincin metalloproteinases, a “cysteine switch” mechanism has been described, based on the presence of a cysteine residue in the pro-segment that prevents access of substrates to the active site of the enzyme and thus keeps it inactive (18). With proteolysis of the pro-segment, the enzyme would become active. A conserved motif surrounding the cysteine residue has been described, which in the case of GP63 from *Leishmania* sp. is HRCIHD or HRCVHD. Using the motif-based sequence analysis tools of the MEME Suite, we examined the cysteine residue at the amino-terminal ends of *T. cruzi* GP63 sequences. Our findings reveal that, except for nearly half the sequences of group G11—where the cysteine is substituted by an arginine—the cysteine residue remains conserved across all other groups where a similar motif was detectable, despite significant variations in the adjacent sequence regions (see Supplementary Table 4). An additional outlier is observed in one sequence from group G2, characterized by a deletion in this specific region.

### 5 Expression profiles of GP63 genes

Previous studies have demonstrated the upregulated transcriptional expression of certain multigene families, such as TS (28) and MASP (2), during the trypomastigote stage (a non-replicative, infective stage interacting with the host). In contrast, the transcriptional profile of GP63 genes remains less defined. In fact, it has been suggested that while one subset of GP63 genes is minimally expressed, another exhibits increased expression in trypomastigotes, and a different subset shows elevated expression in amastigotes (29). However, the reliability of these findings might be compromised, as these analyses were based on collapsed genomes, potentially failing to represent all gene copies accurately.

We have analyzed their expression profiles at the mRNA level with previously generated transcriptomic data from three stages of the parasite (20), observing that each group shows different expression levels and profiles and that several groups and/or members show differential expression at the trypomastigote stage compared to the epimastigote and amastigote stages. As can be seen in Figure 5, while groups G1, G4, G6 and G10 do not show differential expression profiles, the rest of the groups contain genes that are significantly expressed more in the trypomastigote stage compared to the epimastigote and amastigote stages. Remarkably, insertions in the active site of group G3 sequences appear to not affect their expression, as they are distinctively upregulated during the trypomastigote stage, suggesting a functionality for this group. Expression levels were also very different between the groups, with those of group G2 and group G9 being the most highly expressed genes, reaching levels between one and two orders of magnitude higher than the rest of the groups (Figure 5 and Supplementary Table 5). These differential expression profiles do not correlate with the location of genes in the two genomic compartments. Groups showing differential expression are located in both the core and disruptive compartments.

Moreover, the two groups with the highest expression at the RNA level are one in the core and the other in the disruptive compartment. This suggests that differential expression mechanisms are strongly linked to features associated with gene structure, e.g. possible regulatory sequence elements present in the UTRs.

### 6 Patterns of divergence in regions embedding (surrounding) GP63 coding sequences is similar to that of proteins

To determine if the sequence identity observed within each GP63 group extends to untranslated regions (UTRs) or is confined solely to the coding sequence, we conducted a phylogenetic analysis paralleling the approach used for the proteins. This involved analyzing 450 nucleotides upstream (5’ UTR) and 700 nucleotides downstream (3’ UTR) of each GP63 gene, which represent the average length of the UTRs present in the GP63 genes. As illustrated in Figure 6, the clustering patterns in the 5’ and 3’ UTRs largely mirror those seen in the phylogenetic analysis of the proteins. This suggests that GP63 genes within each group not only share sequence identity in their coding regions but also in their UTRs, with the similarity being more pronounced in the 3’ UTR than in the 5’ UTR. As previously discussed, tandem duplication represents a form of gene amplification within this family, with conservation in these regions indicating that duplication encompasses more than just the coding sequence. Moreover, the untranslated regions (UTRs), particularly the 3’UTR, are pivotal in gene expression regulation (30). Hence, the conserved clustering of these sequences might suggest a differential expression regulation, potentially varying across developmental stages or in response to different stress conditions. Notably, in the 3’UTR phylogeny, group G4 undergoes a division, with three sequences diverging from this group (indicated by an arrow in Figure 6). These sequences are the 3’UTRs of genes C4B63_170g41, C4B63_320g17, and C4B63_32g57, which represent the non-expressed genes of G4 at any developmental stage (see Figure 5 and Supplementary Table 5 for reference). Similarly, group G11 bifurcates, with 7 out of the 32 3’UTRs clustering separately. The genes corresponding to these 3’UTRs show very low expression levels during the trypomastigote stage, in contrast to the remaining genes, which exhibit increased expression during this phase.

**Figure 6:**
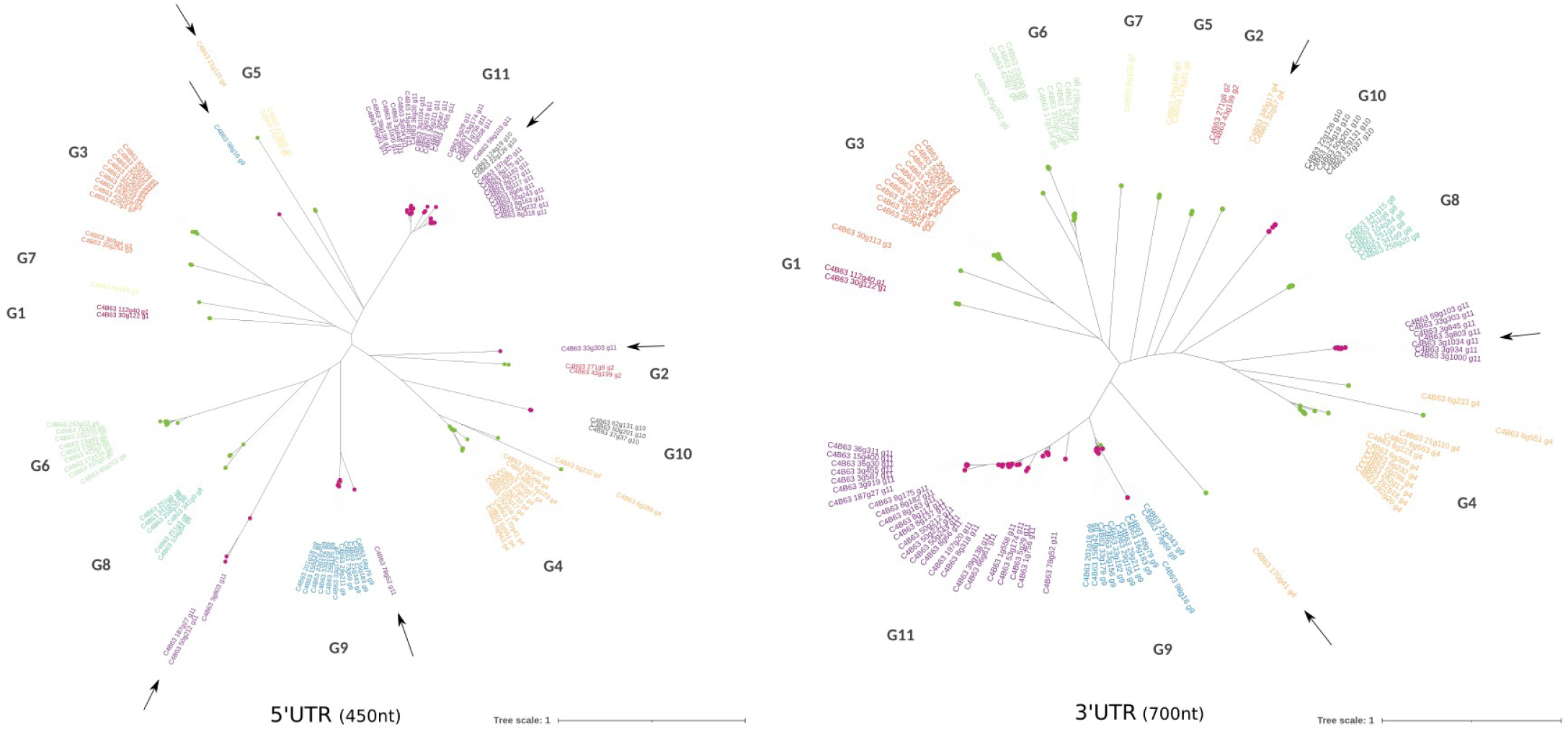
Phylogenetic Analysis of GP63 UTRs. Phylogenetic analysis of the 5’UTRs (450nt) on the left and the 3’UTRs (700nt) on the right. Groups are differentiated by color coding, and labels include the name of the respective gene and group. UTRs that do not cluster with their respective groups are highlighted with a black arrow. Additionally, nodes are colored based on the genomic compartment to which they belong.

Together, all these results suggest that this family of proteins with different levels of mRNA expression may be strongly regulated at the post-transcriptional level by elements present in the UTRs, as has been described for *Leishmania* genes (16)

### 7 Amplification of GP63 genes is partially mirrored in other trypanosomes

The eleven groups described in section 2, most likely reflect functional diversification in the GP63 family of *T. cruzi*. It is of great interest to determine whether this divergence is observed in other species of the genus *Trypanosoma*, other trypanosomatids and even outside the trypanosomatidae family. For this purpose, we conducted a phylogenetic analysis including all completed GP63 sequences identified in eight *Trypanosoma* species, covering the taxonomic diversity of the genus *(Trypanosoma cruzi*, *T. cruzi marinkellei*, *Trypanosoma rangeli*, *Trypanosoma tehilieri*, *Trypanosoma grayi*, and three african trypanosomes). We also included the genus *Strigomonas* (*Blastocrithidia*), two *Leishmania* species and the free living kinetoplastid *Bodo saltans*. The two phylogenetic trees presented in Figure 7 allow us to highlight different aspects of this divergence. These two trees differ in that Figure 7A contains just one representative copy of each group from the large *Trypanosoma* cluster, allowing an easier identification of the divergence outside the genus. From Figure 7A it is evident that ten of the eleven groups described above, represent *Trypanosoma* specific subfamilies, whereas homologues to G1 are observed outside the genus. Interestingly, members of G1 subfamily are found as single copy genes not only in *T. cruzi* but in all species of *Trypanosoma* and *Strigomonas*. The gene is also present in *B. saltans* as multicopy but absent in *Leishmania*. The phylogenetic location of G1, as the earliest branching lineage, and its widespread presence in kinetoplastids is indicative of its ancestrality. This, in turn, is reflected in two aspects of its biology that are worth mentioning. First, it is the only group that lacks the conserved methionine of the Met-turn motif (see Figure 4) and contains a spacing of 71 residues between the conserved Glycine in the active site and the third histidine, which is unique to this group. Second, in *T. cruzi,* the G1 group is transcribed at constant levels throughout the life cycle, suggesting an absent or weak relationship with the infective phase, a feature that is compatible with its presence in the non-infective kinetoplastid *B. saltans*. Analysis of the conserved active site elements in sequences clustered with the *T. cruzi* G1 group revealed the preservation of all elements, including the characteristic substitution of methionine with leucine, which distinguishes this group from others. However, the distance between the active site glycine and the third histidine exhibited some variability, ranging from 69 to 74 amino acids across the sequences. A notable deviation was observed in the *T. congolense* sequence, which contains a deletion resulting in a shorter separation of 58 amino acids between these elements

**Figure 7:**
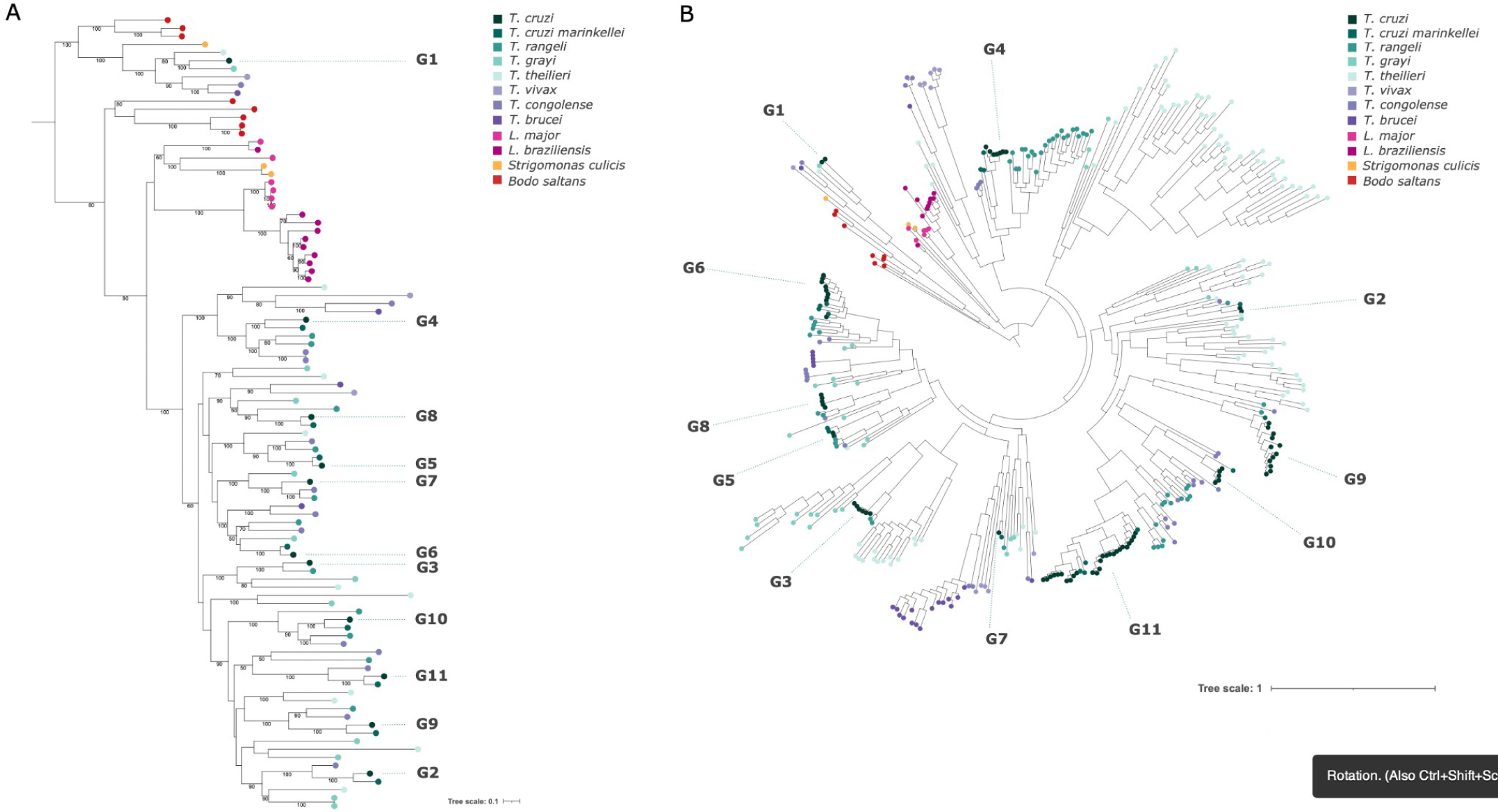
Global Phylogenetic Analysis of GP63 Genes Across Kinetoplastids. A) This panel shows a phylogenetic analysis utilizing the Maximum Likelihood method based on the JTT model with gamma distribution and 100 bootstrap pseudoreplicates. It focuses on the entire GP63 amino acid sequences from representative genes of *T. cruzi, T. cruzi marinkellei, T. rangeli, T. grayi, T. theileri, T. vivax, T. congolense, T. brucei, Leishmania major, L. braziliensis, Strigomonas culicis*, and *Bodo saltans*. Bootstrap support values above 50 are indicated at the respective nodes. B) Presents a phylogenetic analysis of the M8 domain using an enriched dataset of 282 sequences from a diverse array of *Trypanosoma* and *Leishmania* species, including the free-living kinetoplastid *Bodo saltans*, highlighting the evolutionary relationships within this gene family across kinetoplastids.

Another interesting aspect from Figure 7A, is the pattern of amplification and divergence of GP63 in both *L. major* and *L. braziliensis*. The interest in understanding the divergence pattern in these two species and its comparison with *T. cruzi* is fundamentally due to the fact that this is a classical and extensively studied *Leishmania* protein (in fact is called leishmanolysin). Noteworthy, there are only three relatively divergent gene copies, one of which forms a cluster that has 9 copies in *L. braziliensis* and 4 in *L. major* (31). Similar to what is observed in *T. cruzi*, the degree of amino acid divergence among copies within the same cluster is very modest, this being true for both *Leishmania* species included in this study. Besides, similarities are higher within, than among species. This may indicate that the amplifications emerged independently (i.e. are species specific), or that there is a high rate of gene conversion. We note that with this comparatively low copy number and differentiation within cluster in *Leishmania,* this protease is able to exert different functions, facilitating adhesion and invasion and cleaving or degrading several host cell proteins that play important roles in the cellular response, such as transcription factors and components of signaling cascades (9).

Figure 7B is basically an enlargement of part A that incorporates the intragroup variability within the eight species of *Trypanosoma* genus, thus including all sequences with a complete M8 domain (378). As previously noted, ten of the eleven *T. cruzi* groups, defined in section 2, form part of a large clade. A relevant observation is that members of these ten groups are also found in other *Trypanosoma* species, including African ones. There are some exceptions though: group G3 is absent in African trypanosomes, and groups G4 and G9 are not found in *T. grayi* and *T. theilieri*. The copy number of each group exhibits considerable variation across different species, with group G11 notably expanded in *T. cruzi*. *T. cruzi* exhibits very high intra-cluster similarity, in contrast to what is observed in other species of *Trypanosoma* genus (and *Bodo*). This pattern of *T. cruzi* is to some extent similar to that of *Leishmania.* Besides the groups already described in *T. cruzi*, additional large clusters are present in other species with no representation in *T. cruzi*. One of such clusters, designated LTGC1 (Large Theiliri Gp63 Cluster 1), includes over 35 copies in *T. theilieri*, with a few distant relatives in *T. grayi*. A second notable expansion in *T. theilieri*, termed LTGC2, comprises approximately 20 copies. Genes within these non-*T.cruzi* GP63 groups also possess a complete M8 domain and signal peptide, suggesting their likely functionality.

## CONCLUSIONS

In this study we address the amplification and diversification of GP63 genes in the genome of *T. cruzi*, using as an initial reference our previous genomic studies on the Dm28c strain. Our analysis identified the existence of eleven groups, each distinguished by specific characteristics such as sequence length, unique features within the conserved elements of the M8 motif and different expression profiles. Notably, these groups exhibit variations in the presence of signals in the N- and C- terminal regions as well as in the number of sites for N-glycosylation. While there is remarkable homogeneity within each group, substantial divergence is observed when comparing sequences across different groups.

Furthermore, we found a strong correlation between these GP63 groups and the core and disruptive genomic compartments of the *T. cruzi*. Multi-copy tandem arrays of GP63 genes are prevalent in groups situated in the core compartment, whereas those in the disruptive compartment are typically flanked by genes from other multigene families, such as mucins, MASPs, and trans-sialidases.

This study has also allowed us to elucidate the evolutionary expansion of the GP63 gene family in *Trypanosoma.* Phylogenetic analysis provided insights into the evolutionary expansion of the GP63 gene family in *T. cruzi*, showing that this family has followed different evolutionary paths, not only within the genus *Trypanosoma* but also in other species. Ten of the 11 groups characterized in this study are shared only within the genus *Trypanosoma* including African species, while only group G1 is also present in other trypanosomatidae species and nonparasitic kinetoplastids, indicating that group G1 represents the earliest branching lineage. These studies reveal on the one hand, that the expansion of the remaining GP63 groups genes occurred prior to the separation of *Trypanosoma* species. On the other hand, the expansion of GP63 observed in species of the genus *Leishmania* is independent of that of *Trypanosoma*, and vastly more modest.

## METHODS

### GP63 Metalloprotease Sequences

We downloaded 378 sequences annotated as "surface_protease_GP63" within the *T. cruzi* Dm28c genome, from the database http://bioinformatica.fcien.edu.uy/cruzi/. Among them, 96 were identified as genes and 282 as pseudogenes.

To validate and refine these annotations, we focused on sequences that exceeded 460 amino acids in length and possessed a complete M8 peptidase domain, indicative of GP63 genes. This evaluation was performed using the InterPro predictor (32) and CDVist (33), a web server designed to maximize domain coverage in multidomain protein sequences, which enabled the identification of domains within the GP63 protein sequences. N-glycosilation site prediction were done using the NetNGlyc - 1.0 server that predicts N-Glycosylation sites proteins examining the sequence context of N-X-S/T sequences (https://services.healthtech.dtu.dk/services/NetNGlyc-1.0/).

The presence and location of signal peptide cleavage sites was predicted using SignalP 6.0 server (34) selecting Eukaria as an organism. The presence of a glycosylphosphatidylinositol (GPI) anchoring was predicted using two different web-servers: PredGPI (35), and NetGPI 1.1 (https://services.healthtech.dtu.dk/services/NetGPI-1.1/).

### Reannotation of GP63 Pseudogenes

A homology search was performed using blastn, with functional GP63 sequences as a reference database. Additionally, the M8 domain was searched as previously described. Sequences located in close proximity or at short distances were meticulously analyzed using both blastn and blastp against GP63 genes, with YASS (36) facilitating dotplot visualizations. This approach allowed us to identify open reading frames (ORFs) as parts of genes that had undergone pseudogenization. Accordingly, we re-annotated these sequences, extending them from the first nucleotide (nt) of the initial sequence to the last nt of the final constituent sequence, marking the internal ORFs as derived fragments. The "new" parental pseudogene was re-annotated with an ID ending with a letter p.

### Cluster analysis based on amino acid identity

Identity matrix of the GP63 sequences was generated by the pairwise sequence alignment tool Clustal Omega (37) (https://www.ebi.ac.uk/Tools/msa/clustalo/). Visualization of the clustering was achieved with Gephi, where each edge’s color tone represents the degree of identity between sequences. An edge weight threshold of 65% was applied for visualization.

Additionally, a condensed identity matrix, summarizing average intra-group and inter-group identity, was derived from mean group identity analyses. This condensed matrix was visualized using R.

### Gene environment analysis

We analyzed the genomic context of each GP63 gene by identifying the three upstream (5’) and three downstream (3’) genes to assess the gene composition surrounding GP63 loci. For GP63 genes residing in both core and disruptive compartments, we determined the most prevalent gene types, utilizing the annotations of these neighboring genes with a custom script in Perl. The results were plotted using R.

### Expression profile of GP63 genes

Expression profiling for each gene was derived from the work of (20). For each gene, Transcript Per Million (TPM) values were calculated across replicates and developmental stages (amastigote, epimastigote, and trypomastigote), with differentially expressed genes listed in Supplementary Table 5. Rather than using arbitrary TPM thresholds, expression levels were categorized based on quartile distribution of TPM values within each stage: genes in the lowest quartile (<25%) were classified as *low*, those between the 25th and 75th percentiles were deemed *medium*, and genes in the highest quartile (>75%) were considered *high*.

### Phylogenetic analyses

The peptide sequences of the 96 GP63 genes were aligned using MAFFT (38), employing the ‘-einsi’ option for enhanced accuracy in alignments involving sequences with multiple conserved domains and gaps. Subsequent trimming was executed utilizing the ‘gappyout’ option in TrimAl to remove poorly aligned regions and gaps. Phylogenetic analysis was conducted using PhyML (39) with the following parameters: ‘phyml -i 2024_M8.einsi.gappy.phy -b -1 -d aa’. Visualization of the resulting phylogenetic tree was created using iTOL (40).

To enrich the taxonomic coverage of the dataset, 282 sequences from a variety of *Trypanosoma* and *Leishmania* species, and the free living kinetoplastid *Bodo saltans*, including *T. cruzi marinkellei* (17), *T. theileri* (103), *T. grayi* (33), *T. rangeli* (55), *T. congolense* (11), *T. vivax* (12), *T. brucei* (23), *L. major* (6), *L. braziliensis* (10), *Strigomonas culicis* (3), and *Bodo saltans* (9), were incorporated. The M8 domain was identified in these sequences, and both alignment and phylogeny were specifically performed on these extracted domains, as previously mentioned.

From this expanded set of 378 sequences (282 plus 96 from *T. cruzi*), we selected representative sequences for each group of *T. cruzi* GP63 and all other clusters, totaling 103 sequences. The alignment of the selected sequences was done using a MAFFT (38), retaining all positions containing information in at least 80% of the sequences. A phylogeny was then constructed using the Maximum likelihood method using JTT model with gamma distribution and 100 bootstrap pseudoreplicates, providing insights into the evolutionary relationships and diversity within the GP63 gene family across these protozoan organisms.

UTR phylogenies were performed using 450 and 700 nucleotide positions upstream and downstream of 5’ and 3’ ends, respectively. The sequences were retrieved from tritrypdb.org (41), aligned with MAFFT (38) using -auto option, and the phylogenetic tree was constructed with IQ-TREE (42) with default parameters.

## SUPPLEMENTARY FIGURES

**Supplementary Figure 1.**
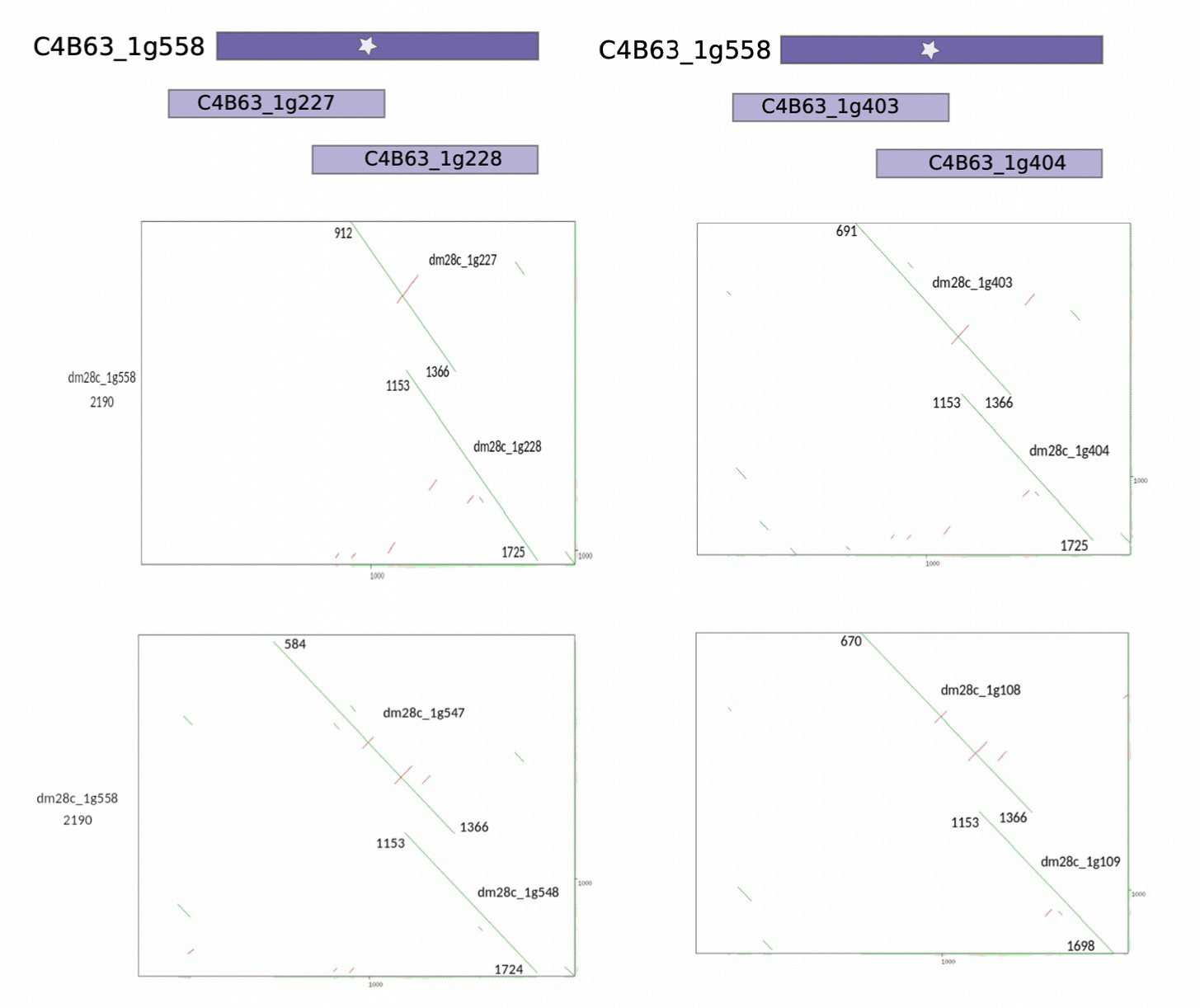
Reassignment of pseudogene fragments, presenting two examples at the top. In these instances, alignments were conducted using the YASS tool, with the C4B63_1g558 gene serving as the reference. Below, the figure provides a visualization of the pairwise local alignments for four pseudogene pairs, demonstrating the consistent pattern observed in the examples above.

**Supplementary Figure 2.**
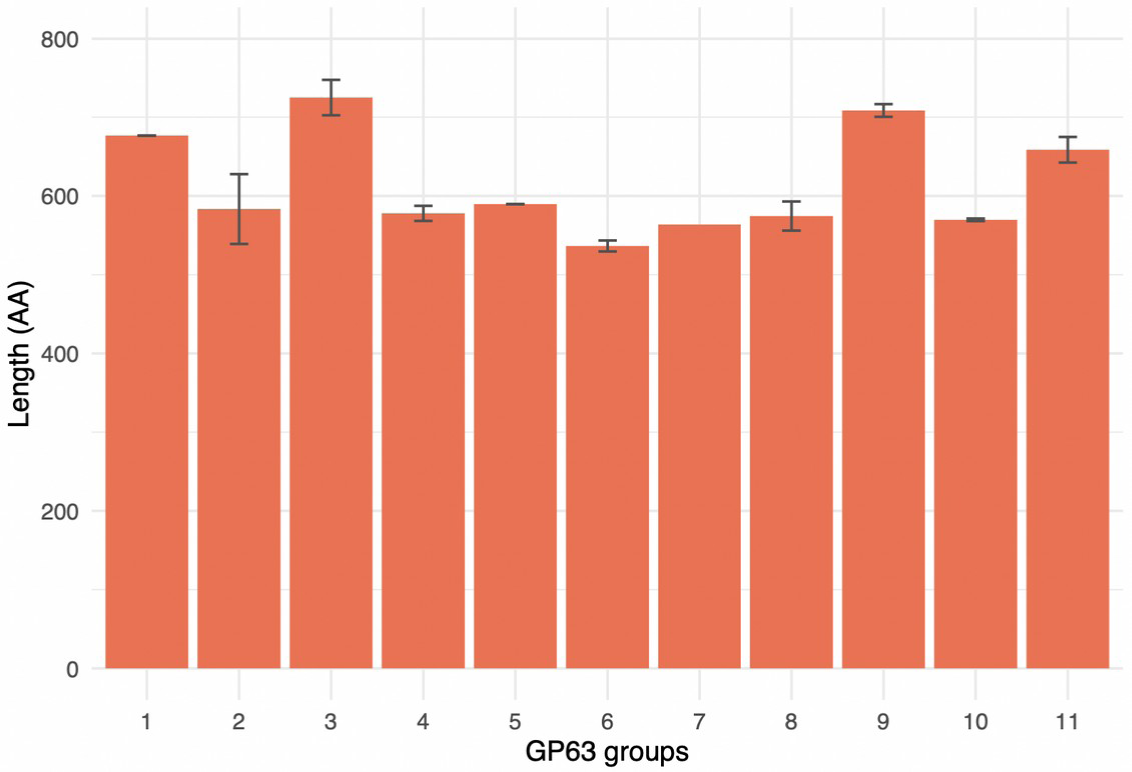
Average length of the amino acid sequences for each GP63 group, including the standard deviation.

**Supplementary Figure 3.**
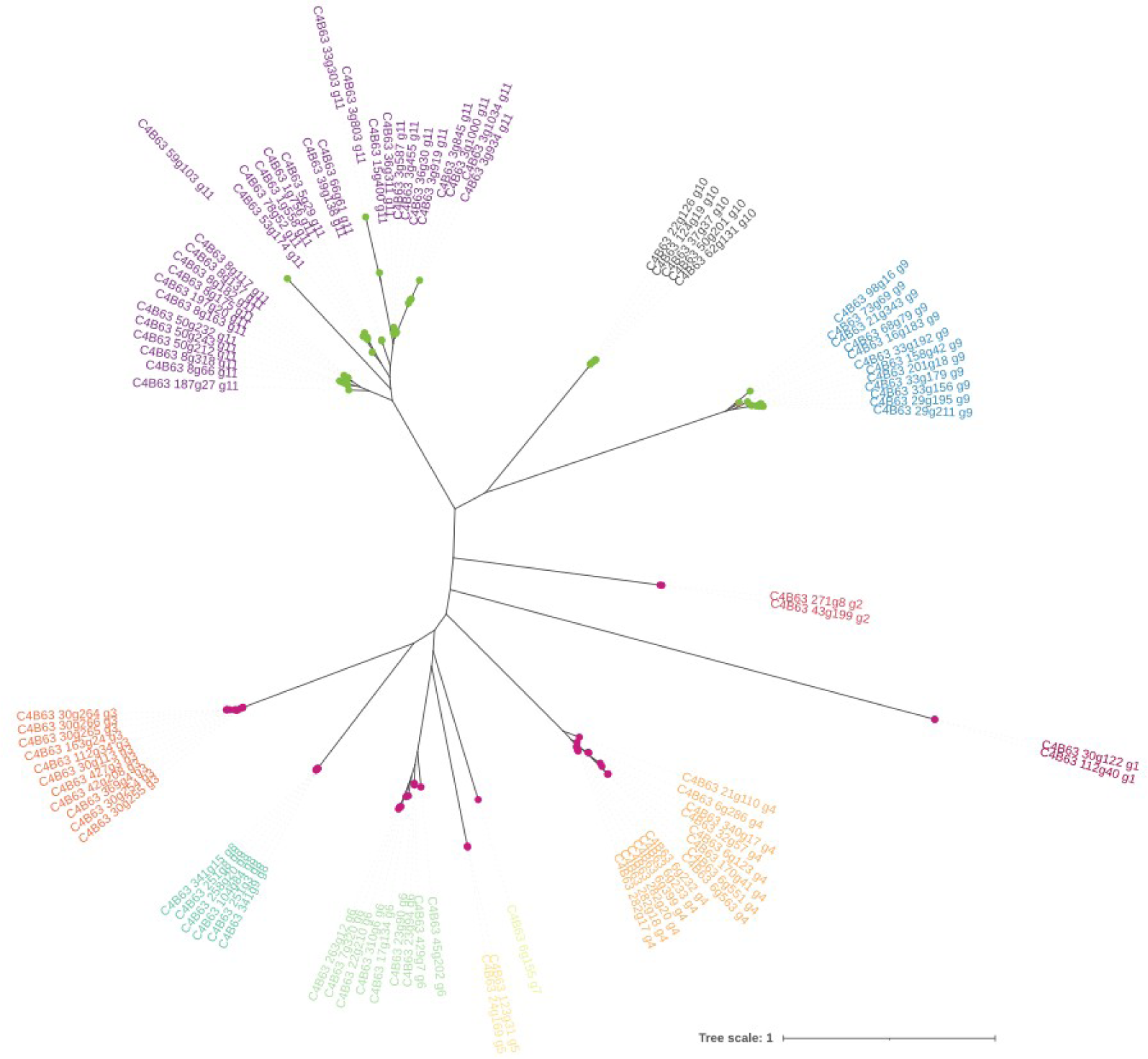
Phylogenetic analysis of the 96 GP63 amino acid sequences from *T. cruzi* Dm28c. Groups are differentiated by color coding in labels. Nodes are colored based on the genomic compartment, core in green, disruptive in purple.

**Supplementary Figure 4.**
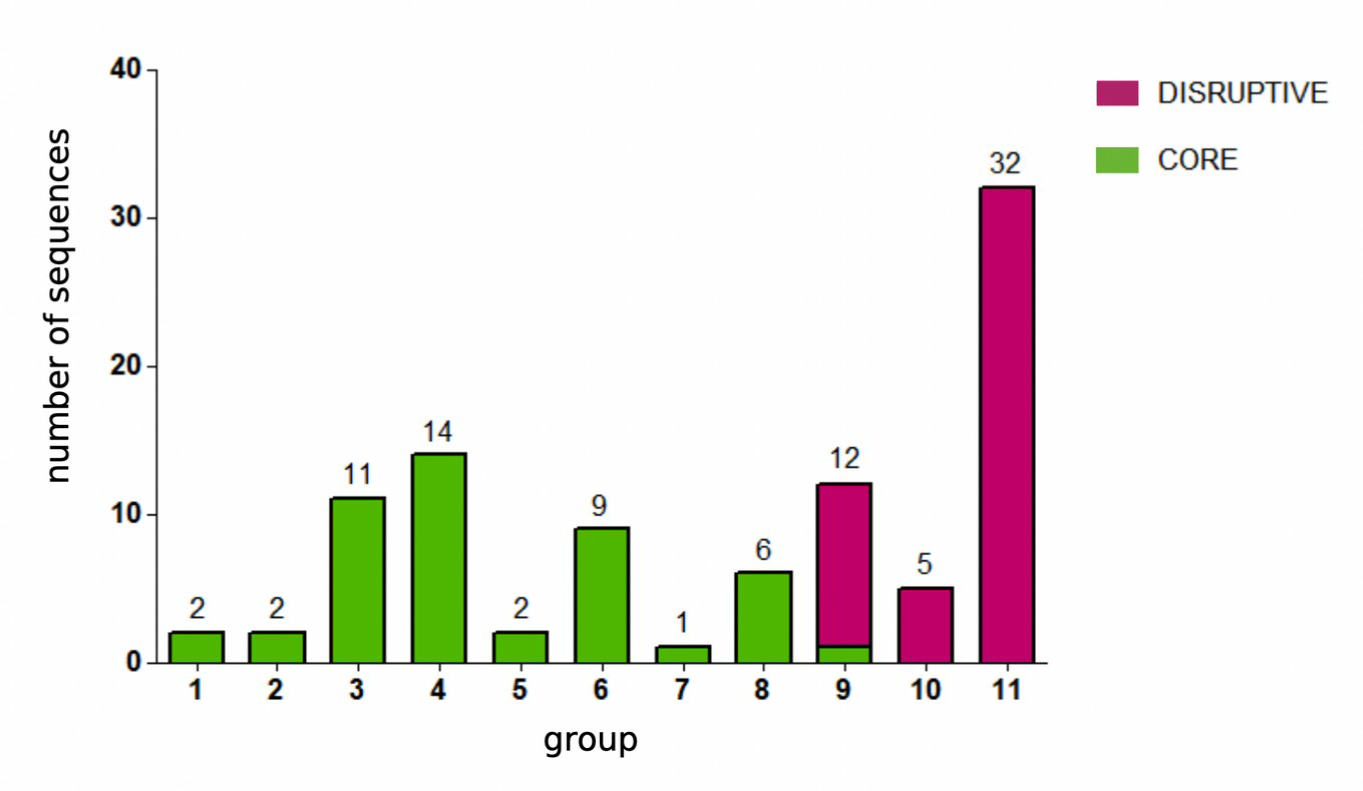
Genomic distribution of GP63 gene sequences across *T. cruzi* groups. Green color corresponds to groups present in the core compartment, purple to groups in the disruptive compartment.

**Supplementary Figure 5.**
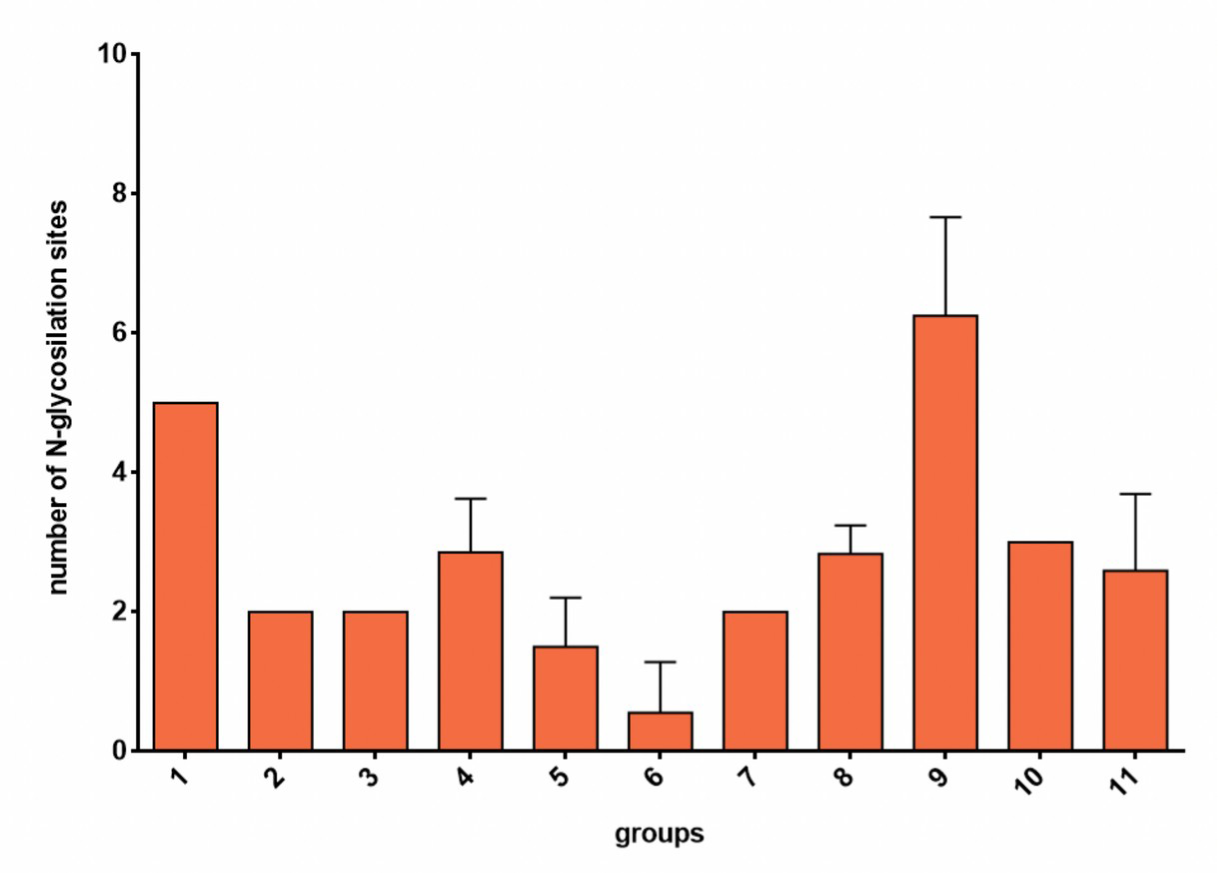
Average number of predicted N-glycosylation sites in each group of GP63 sequences.

## Consent for publication

All authors have approved the manuscript and agree with its submission.

